# Thermal tolerance and preference are both consistent with the clinal distribution of house fly proto-Y chromosomes

**DOI:** 10.1101/2021.01.14.426736

**Authors:** Pablo J Delclos, Kiran Adhikari, Oluwatomi Hassan, Jessica E Cambric, Anna G Matuk, Rebecca I Presley, Jessica Tran, Vyshnika Sriskantharajah, Richard P Meisel

## Abstract

Selection pressures can vary within localized areas and across massive geographical scales. Temperature is one of the best studied ecologically variable abiotic factors that can affect selection pressures across multiple spatial scales. Organisms rely on physiological (thermal tolerance) and behavioral (thermal preference) mechanisms to thermoregulate in response to environmental temperature. In addition, spatial heterogeneity in temperatures can select for local adaptation in thermal tolerance, thermal preference, or both. However, the concordance between thermal tolerance and preference across genotypes and sexes within species and across populations is greatly understudied. The house fly, *Musca domestica*, is a well-suited system to examine how genotype and environment interact to affect thermal tolerance and preference. Across multiple continents, house fly males from higher latitudes tend to carry the male-determining gene on the Y chromosome, whereas those from lower latitudes usually have the male-determiner on the third chromosome. We tested whether these two male-determining chromosomes differentially affect thermal tolerance and preference as predicted by their geographical distributions. We identify effects of genotype and developmental temperature on male thermal tolerance and preference that are concordant with the natural distributions of the chromosomes, suggesting that temperature variation across the species range contributes to the maintenance of the polymorphism. In contrast, female thermal preference is bimodal and largely independent of congener male genotypes. These sexually dimorphic thermal preferences suggest that temperature-dependent mating dynamics within populations could further affect the distribution of the two chromosomes. Together, the differences in thermal tolerance and preference across sexes and male genotypes suggest that different selection pressures may affect the frequencies of the male-determining chromosomes across different spatial scales.

**Impact Statement:** Genetic variation within species can be maintained by environmental factors that vary across the species’ range, creating clinal distributions of alleles responsible for ecologically important traits. Some of the best examples of clinal distributions come from temperature-dependent phenotypes, such as thermal tolerance and preference. Although genotype and developmental temperature strongly affect physiological and behavioral traits in ectotherms, the correlation between these traits across genotypes and sexes within species is greatly understudied. We show that two different male-determining chromosomes found in natural populations of house flies affect both thermal tolerance and preference in a way that is concordant with their clinal distributions across latitudes. This provides strong evidence that temperature variation across the species range contributes to the maintenance of the polymorphism. Furthermore, we find evidence that thermal preference is sexually dimorphic, suggesting that temperature-dependent mating dynamics could further affect the distribution of genetic variation in this system. Therefore, at a macro-geographical scale, the differences in thermal tolerance and preference across male genotypes likely contributes to the maintenance of the cline. Within populations, differences in thermal preference likely affect sexual selection dynamics, which may further affect the frequencies of the chromosomes.

## Introduction

Ecological variation across a species’ range can select for local adaptation within populations, which can contribute to the maintenance of genetic variation by favoring different alleles across the range (Levene, 1953; Felsenstein, 1976; Hedrick *et al.*, 1976; Kawecki & Ebert, 2004). In addition, heterogeneous selection pressures that are distributed as a gradual continuum from one end of the species’ range to another can create a cline of genetic variation responsible for phenotypes under selection (Slatkin, 1973; Endler, 1977). Some of the best examples of latitudinal clines come from temperature-dependent phenotypes (e.g., body size, developmental rate, and thermal tolerance) that have been well-documented in flies (Partridge *et al.*, 1994; Eanes, 1999; Robinson & Partridge, 2001; Hoffmann *et al.*, 2002). Moreover, heterogeneous selection pressures across a cline may affect males and females differently (Connallon, 2015; Connallon *et al.*, 2019), although the empirical evidence for such variation in sex-specific selection across geographic ranges is mixed (Delcourt *et al.*, 2009; Delph *et al.*, 2011; Allen *et al.*, 2017; Lasne *et al.*, 2018).

Thermal adaptation within populations and across a species range can occur via selection on physiological, anatomical, or behavioral traits. For example, north-south gradients in heat and cold tolerance have been observed in *Drosophila* (Hoffmann *et al.*, 2002), suggesting physiological adaptation to thermal environments. In addition, ectotherms, such as flies, rely on behavioral mechanisms of thermoregulation by avoiding suboptimal temperatures in search of more optimal ones (Dillon *et al.*, 2009; Kearney *et al.*, 2009), and thermal preference may be correlated with optimal thermal performance (Dawson, 1975; Angilletta *et al.*, 2002).

Concordance across genotypes between different thermal traits could reinforce the response to selection, whereas negative correlations could constrain adaptation (Etterson & Shaw, 2001). However, it is not clear if physiological and behavioral thermal traits are genetically correlated within a species, between sexes, or across populations (Dawson, 1975; Angilletta *et al.*, 2002; Gilbert & Miles, 2017). For example, experiments in *Drosophila subobscura* identified individual chromosomes that affected thermal tolerance or temperature preference, but no single chromosome affected both physiological and behavioral phenotypes (Dolgova *et al.*, 2010; Rego *et al.*, 2010; Castañeda *et al.*, 2019). Furthermore, temperature-dependent traits can affect assortative mating and male reproductive success (Dolgin *et al.*, 2006; Keller & Seehausen, 2012), suggesting inter-sexual differences in thermoregulation could affect genetic variation within populations via sexual selection. These sex-specific selection pressures could also contribute to the maintenance of genetic variation via inter-sexual conflict or context-dependent selection (Kotiaho *et al.*, 2001; Rostant *et al.*, 2015; Meisel, 2018). Despite the importance of inter-sexual differences, previous work did not test for differences in the genetic correlation of thermal traits between males and females.

We used a sex chromosome polymorphism in the house fly, *Musca domestica*, to investigate the concordance of thermal tolerance and preference across clinally distributed male genotypes. House fly has a polygenic sex determination system, in which a male-determining gene has been mapped to all six chromosomes, some males can carry multiple male-determining chromosomes, and a female-determining allele segregates on one chromosome (McDonald *et al.*, 1978; Inoue & Hiroyoshi, 1986; Dübendorfer *et al.*, 2002; Hediger *et al.*, 2010). The *M. domestica male determiner* (*Mdmd*) gene is most commonly found on either the third chromosome (III^M^) or what was historically referred to as the Y chromosome (Y^M^) (Hamm *et al.*, 2014; Sharma *et al.*, 2017). Both III^M^ and Y^M^ are very young proto-Y chromosomes that are minimally differentiated from their homologous proto-X chromosomes (Meisel *et al.*, 2017; Son *et al.*, 2019; Son & Meisel, 2020). Y^M^ and III^M^ are distributed along latitudinal clines on multiple continents in the Northern Hemisphere (Tomita & Wada, 1989; Hamm *et al.*, 2005; Kozielska *et al.*, 2008). Y^M^ is most frequently found at northern latitudes, and III^M^ is more common at southern latitudes (Figure 1A). This distribution suggests that the Y^M^ chromosome confers higher fitness in colder climates, and, conversely, III^M^ confers higher fitness in hotter climates. Therefore, variation in temperature across the species range may create heterogeneous selection pressures that maintain the proto-Y chromosome cline in house fly. Consistent with this hypothesis, seasonality in temperature is the best predictor of the frequencies of the proto-Y chromosomes across natural populations (Feldmeyer *et al.*, 2008).

**Figure 1.**
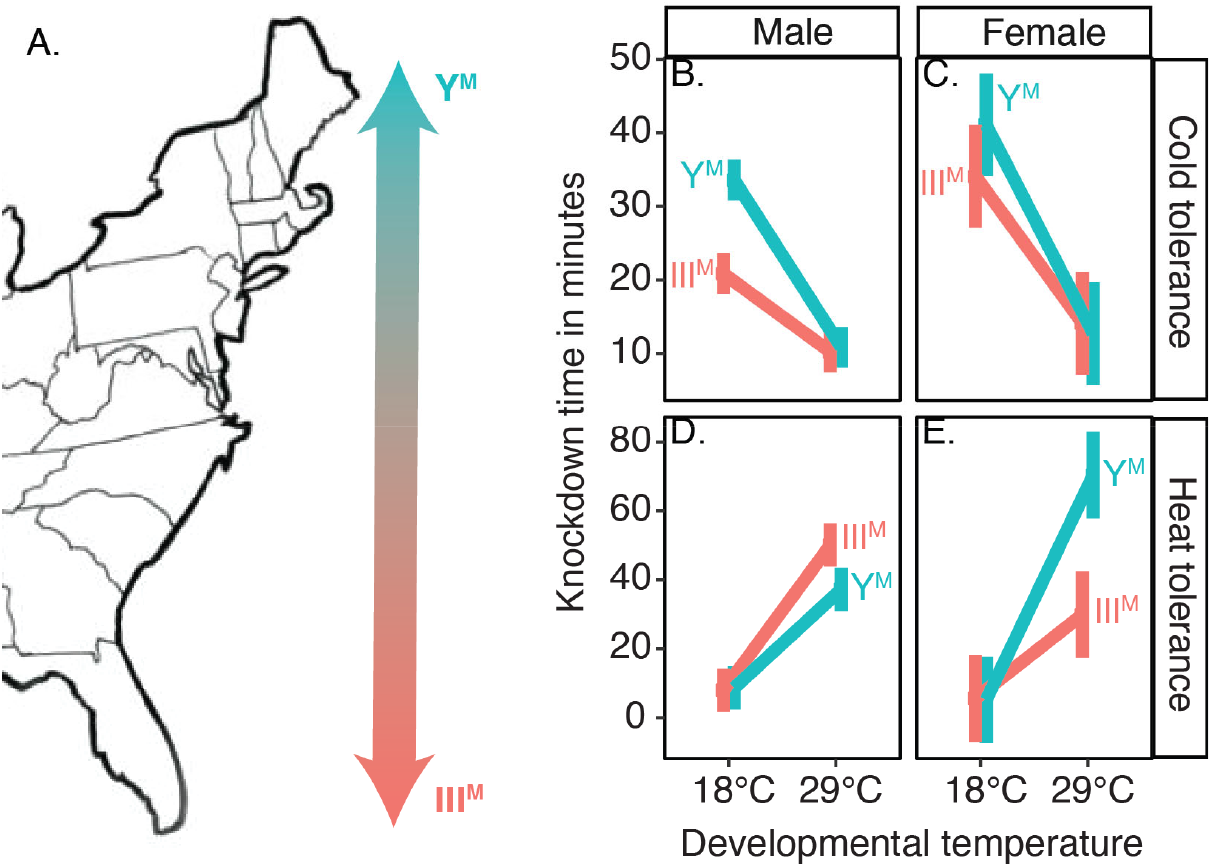
Thermal tolerance in males and females. (**A**) Map of the eastern United States, showing the cline of Y^M^ (more common in the north) and III^M^ (more common in the south). (**B–E**) Graphs show the effect of developmental temperature on knockdown time at either 4°C (cold tolerance) or 53°C (heat tolerance) for Y^M^ (turquoise) and III^M^ (salmon) male flies. Proto-Y chromosome labels for females reflect whether males from the strain carry the Y^M^ or III^M^ chromosome. Mean knockdown time is plotted for each combination of genotype and temperature. Error bars represent standard error.

We tested the hypothesis that the Y^M^ chromosome confers cold-adaptive phenotypes and III^M^ confers heat-adaptive phenotypes in house fly males, which would be consistent with their latitudinal distributions (Tomita & Wada, 1989; Hamm *et al.*, 2005; Feldmeyer *et al.*, 2008; Kozielska *et al.*, 2008). To those ends, we first evaluated if males carrying the III^M^ chromosome (hereafter III^M^ males) have greater tolerance to extreme heat and if males carrying the Y^M^ chromosome (Y^M^ males) have greater cold tolerance. Second, we tested if III^M^ males prefer warmer temperatures than Y^M^ males, and if males and females differ in their thermal preference. We performed all experiments using flies raised at multiple developmental temperatures because thermal acclimation strongly affects temperature-dependent phenotypes in flies and other ectotherms (Krstevska & Hoffmann, 1994; Dillon *et al.*, 2009). Together, we evaluated if thermal preference and tolerance are aligned for sex-linked genetic variants, tested if this alignment is consistent with the geographic distribution of the proto-Y chromosomes, and then discuss how these temperature-dependent phenotypes could affect the access of males to female mates.

## Materials and methods

### Fly strains and rearing

We performed our experiments using five nearly isogenic house fly strains, three with III^M^ males and two with Y^M^ males (Supplementary Methods). All five strains have a common genetic background from an inbred III^M^ strain that was produced from a mixture of flies collected across the United States (Scott *et al.*, 1996; Hamm *et al.*, 2005). Each of the three III^M^ strains carries a different III^M^ chromosome from a separate wild-derived line, and, likewise, the two Y^M^ strains carry different Y^M^ chromosomes. Each strain is fixed for its proto-Y chromosome (either III^M^ or Y^M^), and no other sex determiners, such as the female-determining *Md-tra^D^* allele (Hediger *et al.*, 2010), segregate within these strains.

We reared each strain at 18°C, 22°C, and 29°C for two generations in order to evaluate how thermal acclimation affects thermal tolerance (Chown & Terblanche, 2006) and thermal preference (Krstevska & Hoffmann, 1994; Dillon *et al.*, 2009). Flies from each developmental temperature were assayed at equivalent physiological ages estimated by accumulated degree days (Barnard & Geden, 1993; Wang *et al.*, 2018). For our heat and cold tolerance assays, we used flies 22–50 total degree days after eclosion. For thermal preference assays, we used flies 96–115 total degree days after eclosion. Additional details and calculations are provided in the Supplementary Methods.

### Thermal tolerance

We measured heat and cold tolerance in individual male and female house flies. To measure heat tolerance, lightly anaesthetized individual flies were transferred to a 1.5 ml centrifuge tube that was sealed with fabric. We placed the 1.5 ml tube in a heat block set to 53°C. This temperature was selected because it is the lowest at which heat tolerance could be measured in a reasonable period of time. The time at which a fly fell to the bottom of the tube and could not make its way back to the top was considered the knockdown time. To measure cold tolerance, lightly anaesthetized flies were transferred to a fabric-sealed 20 ml glass vial individually, and the vials were placed in a 4°C refrigerator with a transparent door. Knockdown occured when a fly fell on its back to the bottom of the vial. We gently tapped the assay vial every 2–3 minutes to ensure flies were active.

For both heat and cold tolerance assays, we performed an analysis of variance (ANOVA) using the lmer() function in the lme4 (v1.1) R package (Bates *et al.*, 2015) to model the effect of genotype (G: Y^M^ vs III^M^), developmental temperature (T: 18°C or 29°C), and their interaction on knockdown time (K):

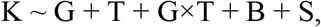

with experimental batch (B) and strain (S) treated as random effects. We also constructed another model excluding the interaction term:

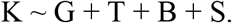

We then used a drop in deviance test to compare the fit of the models with and without the interaction term using the anova() function in R. We also compared heat and cold tolerance between males raised at 22°C and 29°C, using the same approaches as described above. As the thermal tolerance comparisons between flies raised at 18°C and 29°C and between flies raised at 22°C and 29°C were conducted in separate experimental batches, we analyzed each comparison separately. Additional details are provided in the Supplementary Methods.

### Thermal preference

We measured thermal preference as the position of individual flies along a 17–37°C thermal gradient (Figure S1), following a slightly modified version of previous protocols (Anderson *et al.*, 2013; Lynch *et al.*, 2018). For each individual fly, we report mean thermal preference (T_pref_) as the average position during a 10 minute assay window (measured once per minute). We also report thermal breadth, T_breadth_ (Carrascal *et al.*, 2016), as the coefficient of variation of individual-level T_pref_ during the assay window. T_breadth_ provides an estimate of how individuals utilize thermal space within their environment (Slatyer *et al.*, 2013). Choosier individuals show a lower T_breadth_ value and, thus, would be expected to occupy a narrower range of temperatures within a given thermal habitat.

To determine the effects of developmental temperature (18°C, 22°C, and 29°C), genotype (Y^M^ and III^M^), and their interaction on mean T_pref_ across sexes, we created a mixed-effects model using the lme4 package (v1.1) in R (Bates *et al.*, 2015). Developmental temperature, genotype, and their interaction were included as fixed effects, and strain, batch, and lane in the thermal gradient (L) were included as random effects:

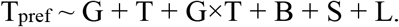

We did the same for T_breadth_. We then determined whether groups significantly differed in T_pref_ or T_breadth_ using Tukey contrasts with the multcomp package (v1.4) in R (Hothorn *et al.*, 2008). Within developmental temperature treatments, we used Bayesian information criterion (BIC) scores from the mclust (v5.4.5) package in R (Scrucca *et al.*, 2016) to determine whether the distribution of individual measures of T_pref_ within a group are best explained by one or multiple normal distributions.

## Results

### Thermal tolerance depends on developmental temperature and male genotype

We measured extreme heat (53°C) and cold (4°C) tolerance as a readout of differences in physiological thermal adaptation between Y^M^ and III^M^ house fly males. We observed the expected effect of acclimation on both heat and cold tolerance (Chown & Terblanche, 2006): flies raised at 18°C tolerate cold longer than the flies raised at 29°C, and flies raised at 29°C tolerate heat longer than flies raised at 18°C (Figure 1). We also find that Y^M^ males are more cold tolerant, and III^M^ males are more heat tolerant, consistent with the latitudinal distributions of Y^M^ and III^M^ males in nature (Tomita & Wada, 1989; Hamm *et al.*, 2005; Feldmeyer *et al.*, 2008; Kozielska *et al.*, 2008). However, the effect of genotype on thermal tolerance depends on acclimation temperature. Specifically, a linear model with an interaction between genotype (Y^M^ or III^M^) and developmental temperature fits the cold tolerance data significantly better than a model without the interaction term (χ²_1_ = 19.3, *p* = 1.1 × 10^−5^). This provides evidence for a G×T effect on cold tolerance—Y^M^ males are more cold tolerant than III^M^ males, but only if they are raised at 18°C (Figure 1B). There is also a significant G×T interaction affecting heat tolerance (χ²_1_ = 4.71, *p* = 0.030 comparing models with and without the interaction term): III^M^ males are more heat tolerant than Y^M^ males, but only if raised at 29°C (Figure 1D).

We next attempted to identify a threshold temperature for the genotype-specific benefits of acclimation by comparing heat and cold tolerance of flies raised at 22°C and 29°C (instead of 18°C and 29°C). We did not observe a significant effect of the interaction between developmental temperature and male genotype on extreme cold tolerance (χ²_1_ = 0.947, *p* = 0.331 comparing models with and without an interaction term) (Figure S2). We therefore hypothesize that there is a threshold temperature between 18°C and 22°C, below which Y^M^ males experience a greater benefit of cold acclimation than III^M^ males. In contrast, there is a significant interaction between genotype and developmental temperature on heat tolerance when comparing males raised at 22°C and 29°C (χ²_1_ = 11.02, *p* = 9.0 × 10^−4^ comparing models with and without the interaction term) (Figure S2). Therefore, the threshold for a genotype-specific benefit from heat acclimation lies between 22°C and 29°C.

We do not expect any difference in heat or cold tolerance across females from our different strains because all females have the same genotype, regardless of the male genotype in the strain. Indeed, a model with an interaction between developmental temperature and male genotype does not fit the female cold tolerance data better than a model without the interaction term (χ²_1_ = 1.46, *p* = 0.23) (Figure 1C). There is a significant effect of developmental temperature on cold tolerance in females (χ²_1_ = 43.5, *p* = 4.3 × 10^−11^ comparing a model with and without developmental temperature), demonstrating that females benefit from cold acclimation regardless of male genotype (Figure 1C). Surprisingly, there is a significant interaction between male genotype and developmental temperature on heat tolerance in females (χ²_1_ = 10.4, *p* = 0.0013 comparing a model with and without the interaction term). In general, females raised at warmer temperatures are more heat tolerant (Figure 1E). However, the interaction of male genotype and developmental temperature is in the opposite direction from what would be expected based on the latitudinal distribution of Y^M^ and III^M^: females from strains with Y^M^ males that are raised at 29°C are more heat tolerant than females from III^M^ strains raised at 29°C (Figure 1E). We thus conclude that the heat and cold tolerance differences between Y^M^ and III^M^ males are specific to males and/or the proto-Y chromosomes (i.e., not genetic background) because we do not observe the same heat or cold tolerance differences in females from those strains (who do not carry the proto-Y chromosomes).

### Thermal preference depends on developmental temperature and male genotype

We next tested if genotype and developmental temperature affect thermal preference (T_pref_). First, we find that T_pref_ is inversely proportional to developmental temperature (Figure 2), with house flies that develop at a warmer temperature preferring cooler temperatures (and *vice versa*), regardless of sex (male: F_2, 742.7_ = 138.4, *p* < 1.0 × 10^−5^; female: F_2, 245.3_ = 37.1, *p* = 1.19 × 10^−4^; Figure 2). This is consistent with how developmental acclimation affects T_pref_ in *Drosophila* (Dillon *et al.*, 2009).

**Figure 2.**
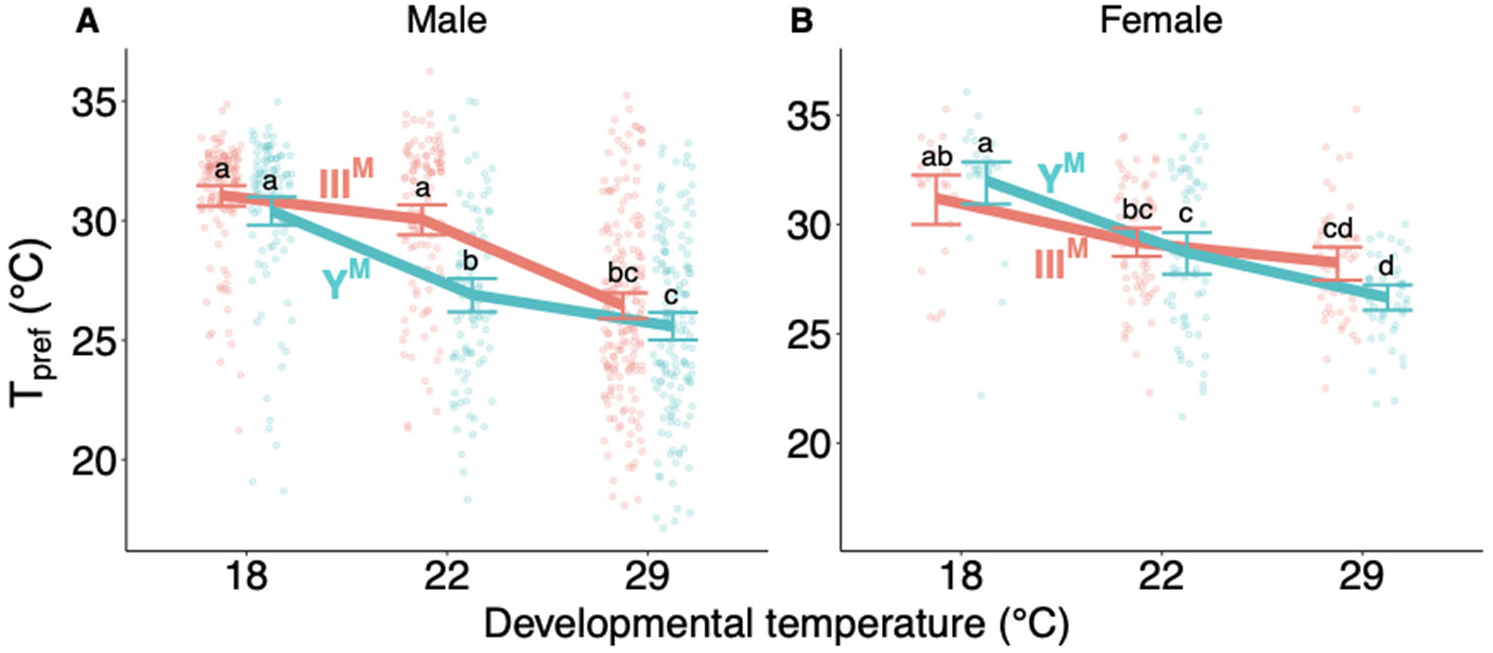
Thermal preference (T_pref_) of (**A**) male and (**B**) female house flies according to male genotype (III^M^ = salmon points and line, Y^M^ = turquoise points and line) and developmental temperature. Each point depicts the mean thermal preference for an individual fly, with lines and error bars denoting means within groups and standard errors of the mean, respectively. Significant differences between groups are denoted by letters, with differing letters highlighting significantly different mean thermal preferences within each graph (Tukey’s *post hoc* test, *p* < 0.05).

We also find that male proto-Y chromosome genotype (Y^M^ vs III^M^) affects T_pref_ (F_1, 756.2_= 44.5, *p* < 1.0 × 10^−5^). There is also a significant interaction effect between developmental temperature and genotype on T_pref_ in males (F_2, 756.3_ = 8.47, *p* = 2.31 × 10^−4^, Figure 2A). Male T_pref_ is similar across genotypes when they develop at either 18°C or 29°C. However, when reared at 22°C, III^M^ males prefer warmer temperatures than Y^M^ males (Tukey’s *post hoc* test, *p* < 0.001). This is consistent with III^M^ males being more common at lower latitudes (where average temperatures are warmer), and Y^M^ males more common at higher latitudes (Tomita & Wada, 1989; Hamm *et al.*, 2005; Feldmeyer *et al.*, 2008; Kozielska *et al.*, 2008). We do not expect differences in T_pref_ in females across strains because all females have the same genotype. Indeed, the genotype of males in a strain (Y^M^ vs III^M^) and the interaction between male genotype and female developmental temperature showed no significant effect on T_pref_ in females (ANOVA, all *p* > 0.1 in Figure 2B). We assayed more males than females in our thermal preference experiments, and so we repeated our analysis by down-sampling the data to have equal numbers of individuals across treatments. The down-sampled data give equivalent results to the full data set (Supplementary Results).

### Thermal breadth depends on sex and thermal preference

We used thermal breadth (T_breadth_) as a measure of the specificity of T_pref_. Male T_breadth_ was not significantly affected by either developmental temperature, genotype, or the interaction between genotype and developmental temperature (ANOVA, all *p* > 0.1, Figure S3A). In contrast, developmental temperature (F_2, 236.9_ = 16.5 , *p* < 1.0 × 10^−5^), as well as the interaction between developmental temperature and male genotype (F_2, 243.6_ = 5.35, *p* = 0.005), had significant effects on T_breadth_ in females. However, the significant interaction is of small effect, as females from strains with differing male genotypes do not significantly differ in T_breadth_ within any developmental temperature treatment (Figure S3B).

The effect of developmental temperature on T_breadth_ in females is driven by increased variance in T_pref_ when females develop at 22°C. The increased variance in female T_pref_ can be explained by a mixture of two normal distributions (Figure 3A, see Table S1 for statistics). This bimodal distribution is not a result of differences across strains because the same pattern was observed among females separately analyzed based on male genotype (Figure S4). In comparison, a single normal distribution best fit Y^M^ male T_pref_ when developed at 22°C, and two normal distributions best explained the III^M^ male T_pref_ when developed at 22°C. Upon inspection, however, the two distributions representing III^M^ male T_pref_ likely correspond to the tail (mean of 28.7°C and large variance of 10.4°C) and peak (mean of 32.6°C and small variance of 0.4°C) of a single skewed distribution, which we are unable to detect using the mclust package we used to fit distributions to our data.

**Figure 3.**
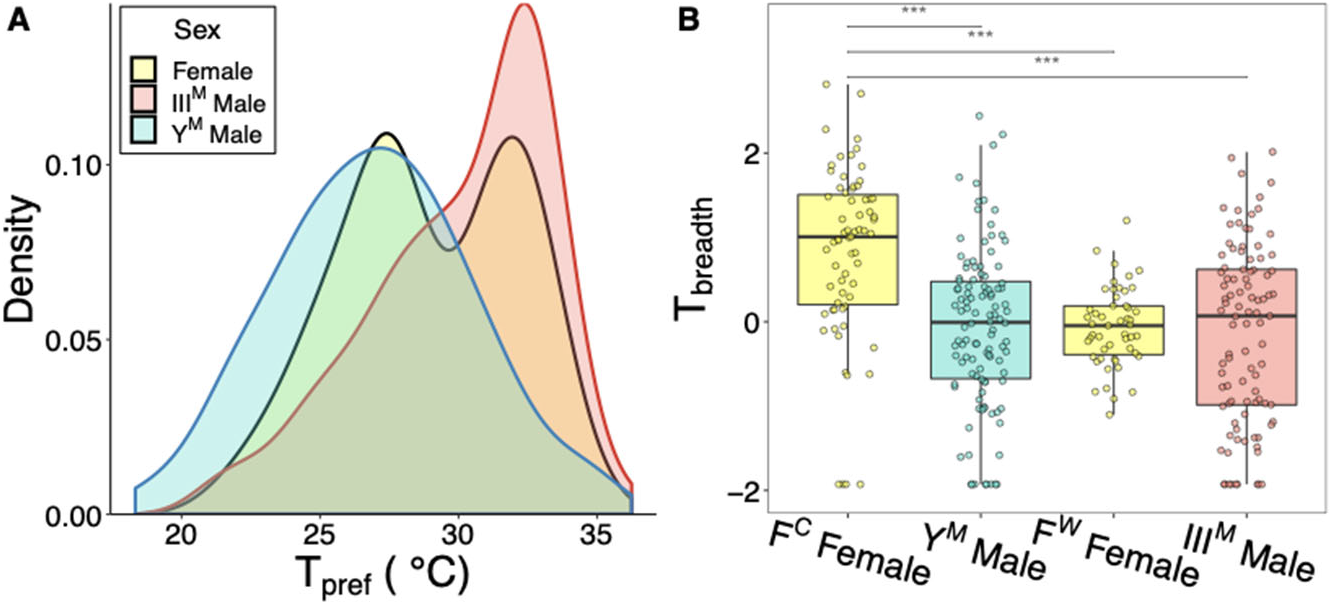
Thermal breadth (T_breadth_) depends on male genotype and sex. (**A**) Distribution of individual-level mean thermal preferences (T_pref_) of III^M^ males, Y^M^ males, and pooled females that developed at 22℃. Y-axis represents relative density of data points and is analogous to frequency of data points for a given T_pref_ value. (**B**) T_breadth_ of individuals raised at 22℃ according to group (F^C^ = cold-preferring females, F^W^ = warm-preferring females). Boxplots denote median values and lower- and upper- quartiles. Asterisks denote significant differences in T_breadth_ between groups (***: Tukey’s *post hoc* test, *p* < 0.01).

We used our model-based clustering analysis of T_pref_ to classify individuals that developed at 22°C into one of four groups: Y^M^ males (lower T_pref_), III^M^ males (higher T_pref_), females with cooler T_pref_ (F^C^ females, 59.3% of females tested), and females with warmer T_pref_ (F^W^ females, 40.7% of females tested). The mean T_pref_ of F^C^ females (26.90°C) is nearly equal to the mean T_pref_ of Y^M^ males (26.87°C; Figure 3A). Similarly, the mean T_pref_ of F^W^ females (32.2°C) is near the mode of the T_pref_ of III^M^ males (32.0–32.5°C; Figure 3A).

We further find that T_pref_ is predictive of T_breadth_ for flies that develop at 22°C. We considered flies from our four T_pref_ groups (Y^M^ males, III^M^ males, F^C^ females, and F^W^ females), and we found a significant effect of group on T_breadth_ (*F*_3, 32.9_ = 9.40, *p* = 1.24 × 10^−4^). Specifically, F^C^ females have significantly greater T_breadth_ than all other groups (Tukey’s *post hoc* test, all *p* < 1.0 × 10^−5^, Figure 3B). Therefore, if we consider T_breadth_ as a measure of the strength of T_pref_, adult house flies can be summarized by one of three phenotypes related to thermal behavior when developed at 22°C: a relatively strong preference for warm temperatures (III^M^ males and F^W^ females, which have high T_pref_ and low T_breadth_), a strong preference for cooler temperatures (Y^M^ males, with low T_pref_ and low T_breadth_), and a relatively weak preference for cooler temperatures (F^C^ females, with low T_pref_ and high T_breadth_). Down-sampling the data gives similar results as the full data set (Supplementary Material).

## Discussion

We tested if thermal tolerance and preference depend on sex and male genotype in house flies. We find that males carrying the Y^M^ chromosome (which is common in the northern end of the species’ range) are more cold tolerant and prefer colder temperatures. Conversely, males carrying the III^M^ chromosome (which is common in the southern end of the species’ range) are more heat tolerant and prefer warmer temperatures. Our results are therefore consistent with the general trend that temperate populations are typically more cold-tolerant than (sub-) tropical ones (Gibert & Huey, 2001; Hoffmann *et al.*, 2002). The differences in thermal preference are consistent with the idea that behavioral thermoregulation can weaken selection for thermal tolerance, as predicted by the “Bogert Effect” (Huey *et al.*, 2003; Huey & Pascual, 2009; Castañeda *et al.*, 2013). However, the fact that thermal preference and tolerance are both predicted by male genotype provides evidence that these traits are responsive to selection, suggesting any Bogert effects are not sufficient to overwhelm thermal adaptation. These differences in thermal tolerance and preference in males depend on developmental temperature, and they are not observed in congener females from the same strains (who do not carry the Y^M^ or III^M^ chromosome). However, females exhibit a bimodal T_pref_, with females from each of the two subgroups overlapping with one of the male genotypes.

### Thermal tolerance and preference depend on developmental temperature, genotype, and sex

Our results demonstrate, to the best of our knowledge, the first documented example of concordant temperature preference, cold tolerance, and heat tolerance across genotypes within a species. We find that Y^M^ males both have greater cold tolerance and prefer colder temperatures, whereas III^M^ males have greater heat tolerance and prefer warmer temperatures (Figures 1 and 2), consistent with their latitudinal distributions (Tomita & Wada, 1989; Hamm *et al.*, 2005; Feldmeyer *et al.*, 2008; Kozielska *et al.*, 2008). Previous work has identified concordant T_pref_ and heat tolerance differences across species (Qu *et al.*, 2011), or found no clear relationship between thermal tolerance and preference across genotypes within species (Yang *et al.*, 2008; Rego *et al.*, 2010; Castañeda *et al.*, 2019). Body size is also predicted to vary with thermal traits (Leiva *et al.*, 2019). In our study, we did not measure insect body size. While we did not observe any obvious differences between strains, it is possible that some of the genotypic effect on thermal tolerance or preference we observed is due to (temperature-dependent) morphological differences between Y^M^ and III^M^ males. Future studies should directly test this hypothesis.

We observed strong effects of developmental temperature on both thermal tolerance and preference that depend on both genotype and sex. Acclimation effects on heat and cold tolerance (Figure 1) are well-documented for ectotherms, including flies and other insects (Bowler & Terblanche, 2008). An inverse relationship between developmental temperature and thermal preference has also been observed in other flies (Dillon *et al.*, 2009; Castañeda *et al.*, 2013). Behaviorally navigating towards compensatory temperatures could serve as a means of mitigating the costs of thermally suboptimal development (i.e., too hot or too cold). The observed relationships between thermal tolerance and developmental temperatures are likely to be caused by acclimation and unlikely to be the result of natural selection within our experiment for two reasons. First, there is unlikely to be sufficient genetic variation in these inbred strains for selection to generate these results within 2 generations. Second, prior attempts at selecting for thermal tolerance in house flies resulted in negligible differences in tolerance across developmental temperatures (Geden *et al.*, 2019) However, it is worth noting that the males used by Geden et al. (2019) were likely all III^M^ based on their geographic origin. Had the experimental population consisted of both III^M^ and Y^M^ males, a response to tolerance may have been detected. We conclude that the differences in thermal tolerance (and preference) between Y^M^ and III^M^ males have evolved across the natural populations from which we sampled the Y^M^ and III^M^ chromosomes.

There are important methodological implications for our observation that variation in thermal preference across genotypes depends on developmental temperature. We only observe warmer (colder) thermal preferences in III^M^ (Y^M^) males when developed at 22°C; thermal preference did not differ between male genotypes when raised at more extreme (18°C, 29°C) temperatures (Figure 2A). Previous studies attempting to estimate genetic variance in thermal preference within or among populations of *Drosophila* have had mixed results. While some studies identified genetic variance among populations within species (Good, 1993; Castañeda *et al.*, 2013), others did not detect substantial variance within (Krstevska & Hoffmann, 1994) or among species (MacLean *et al.*, 2019). Our results show that the phenotypic presentation of genetic variation for thermal preference can depend on the environmental conditions experienced, which could explain why this variance was not detected in other experiments. In addition, while the genetic mechanisms that regulate thermal tolerance in other systems have been extensively studied (Svetec *et al.*, 2011; Königer & Grath, 2018; Königer *et al.*, 2019), it is possible that some of the molecular pathways involved will only be revealed through experiments conducted across developmental temperatures.

We identify multiple differences between males and females in their thermal tolerance and preferences. The strain differences we observed are primarily limited to males, which is expected because the males differ in genotypes (Y^M^ and III^M^) but females are isogenic (Meisel *et al.*, 2015). However, there is a difference in heat tolerance between females from strains with Y^M^ males and females from strains with III^M^ males (Figure 1). While we can rule out certain genotypic explanations for this difference (i.e., all females are isogenic and do not carry *Md-tra^D^*), we do not yet have a mechanistic explanation on why females show the opposite developmental heat tolerance from males. Nevertheless, the difference in heat tolerance observed between females from different strains is in the opposite direction as between Y^M^ and III^M^ males from those strains. This helps us to conclude that differences between Y^M^ and III^M^ males are indeed a result of different proto-Y chromosomes rather than their genetic backgrounds. In other words, the difference in heat tolerance between females is an exception that proves the rule with respect to the effects of proto-Y chromosomes on male thermal tolerance and preference.

We identified a female-specific plasticity for thermal preference that does not map to male genotype. In females, we found that neither thermal tolerance nor thermal preference differ predictably between strains where males carry different proto-Y chromosomes (Figures 1C, E and 2B). However, there is a bimodal thermal preference for females that develop at 22℃ (Figure 3A), regardless of congener male genotype. In addition, females that had colder T_pref_ when developed at 22°C also had a larger T_breadth_ (Figure 3B). In small ectotherms with little thermal inertia, measures of movement along a thermal gradient (such as T_breadth_) are predicted to be positively correlated with environmental temperature (Anderson *et al.*, 2007). However, we observe the opposite relationship between mean environmental temperature (T_pref_) and T_breadth_ in females (Figure 3), suggesting that the difference in T_breadth_ cannot be explained by thermal inertia. Our results suggest that, in nature, females with colder temperature preferences may occupy a wider range of temperatures than females with warmer temperature preferences. Because all females in our experiment are expected to have the same genotype, we hypothesize that these differences in T_pref_ and T_breadth_ are conferred by a plastic response to some yet to be characterized factor (e.g., microclimates within larval rearing containers). Alternatively, this plasticity could have a stochastic origin that is intrinsic to the development of thermal preference (Honegger & de Bivort, 2018; Jensen, 2018).

The correlation between thermal preference and thermal breadth at 22℃ is female-specific: Y^M^ and III^M^ males have similar T_breadth_ values when raised at 22℃ despite their differences in T_pref_. Although general sex differences in thermal tolerance (Hoffmann et al. 2005) and thermal preference (Krstevska and Hoffmann 1994) have been documented, this is the first study, to our knowledge, to identify sex differences in the relationship between thermal preference and thermal breadth. Our results suggest that male and female house flies exhibit different thermoregulatory behavioral patterns which may further be influenced by genotype. Directly identifying a sex-by-genotype-by-environment interaction is beyond the scope of this study because sex and genotype are confounded in our experimental design (the females in our experiment have a different genotype from either male, characterized by a lack of either the III^M^ or Y^M^ chromosome). Nonetheless, the house fly is a tractable system for directly testing for sex-specific genotype-by-environment interactions on thermoregulation. For example, future work could test for sex-specific effects of Y^M^ and III^M^ by measuring phenotypes in females carrying a proto-Y chromosome along with the epistatic female-determining *Md-tra^D^* allele (Hediger *et al.*, 2010; Hamm *et al.*, 2014).

### Environmental heterogeneity and the maintenance of polygenic sex determination

Sex determination pathways rapidly diverge across species, driving evolutionary turnover of sex chromosomes (Bull, 1983; Beukeboom & Perrin, 2014). Polygenic sex determination systems, in which more than one master sex determining locus segregate independently on different chromosomes, have been observed in multiple animal species (Moore & Roberts, 2013). Most population genetic models that attempt to explain the stable maintenance of polygenic sex determination focus on sexually antagonstic effects of sex determining loci or linked alleles on sex chromosomes (Rice, 1986; van Doorn & Kirkpatrick, 2007; Kozielska *et al.*, 2010; Meisel *et al.*, 2016). Less attention has been given to ecological factors that can maintain polygenic sex determination (Pen *et al.*, 2010; cf. Bateman & Anholt, 2017).

Our results demonstrate how spatially variable ecological factors can maintain polygenic sex determination. Specifically, thermal tolerance and preference phenotypes conferred by the Y^M^ and III^M^ chromosomes (Figures 1 and 2) are consistent with the clinal and temperature-dependent distributions of the Y^M^ and III^M^ chromosomes (Tomita & Wada, 1989; Hamm *et al.*, 2005; Feldmeyer *et al.*, 2008; Kozielska *et al.*, 2008). Previous experiments identified multiple fitness advantages conferred by the III^M^ chromosome over Y^M^ at warmer temperatures, including an increase in frequency of III^M^ over generations in a laboratory population (Hamm *et al.*, 2009). However, these fitness differences can only explain the invasion or fixation of the III^M^ chromosome, not the maintenance of the polymorphism. In contrast, differences in thermal tolerance and preference could maintain proto-Y chromosome polymorphism across the species’ range, similar to how selection maintains other clinal variation (Slatkin, 1973; Endler, 1977).

The house fly system reveals how temperature variation can contribute to the maintenance of polygenic sex determination independently of selection on the sex-determination pathway itself. Temperature is an important contributor to the evolution of sex determination pathways in vertebrates (Bull & Vogt, 1979; Holleley *et al.*, 2015). However, the effects of the house fly proto-Y chromosomes on thermal tolerance and preference likely act independently of the sex determination pathway because there are not differences in the expression of sex determination genes across house fly male genotypes raised at different temperatures in a way that is consistent with their clinal distribution (Adhikari *et al.*, n.d.). This suggests that the effects of the Y^M^ and III^M^ chromosomes on thermal phenotypes is a result of alleles on proto-Y chromosomes that are genetically linked to the male-determining locus, as opposed to the male-determiner itself. Therefore, our results highlight how temperature can be important for the evolution of sex determination independently of temperature-dependent activity of the sex determination pathway. Future theoretical work should consider the effect of spatially heterogeneous selection pressures on the maintenance of polygenic sex determination, similar to how temporal heterogeneity can create fluctuating selection pressures that maintain polygenic sex determination (Bateman & Anholt, 2017).

### Selection on thermal phenotypes may depend on geographical scale

Our results suggest that selection on thermal traits differs between macro-geographic species ranges and at a micro-geographical scale within populations. Similar differences in selection pressures according to geographic scale have been documented before in other species (Richter-Boix *et al.*, 2010; De Block *et al.*, 2013; Tüzün *et al.*, 2017). Thermal tolerance and preference in male house flies depend on proto-Y chromosomes genotype in a way that is consistent with the latitudinal distribution of the Y^M^ and III^M^ chromosomes (Figures 1 and 2). This suggests that, at the macro-geographic scale, selection is operating on male physiology and behavior to create or maintain the clinal distribution of Y^M^ and III^M^ (Figure 4A). It is also worth noting that our study focuses on only two male genotypes (III^M^ and Y^M^). While these are the most prevalent genotypes in the eastern United States, other genotypes exist (including males with multiple proto-Y chromosomes, and females with proto-Y and proto-W chromosomes) and are common in other populations (Franco *et al.*, 1982; Feldmeyer *et al.*, 2008; Hamm & Scott, 2009; Hamm *et al.*, 2014). Future studies should characterize thermal tolerance and preference of these other genotypes in order to determine whether their geographical distribution is similarly explained by thermal biology.

**Figure 4.**
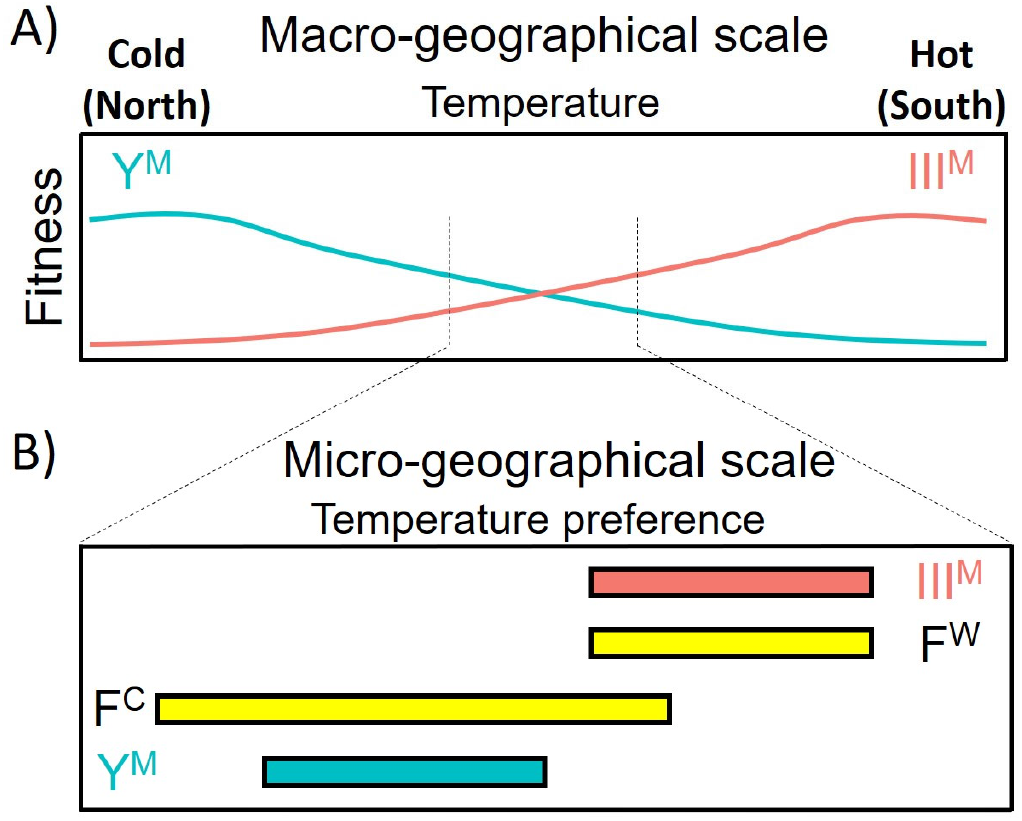
Selection on the III^M^ and Y^M^ chromosomes likely differs across geographic scales. (**A**) At the macro-geographical scale, selection for thermal tolerance and/or thermal preference results in the clinal distribution of the Y^M^ (turquoise) and III^M^ (salmon) chromosomes. (**B**) At intermediate developmental temperatures, male genotypes (Y^M^ vs III^M^) differ in thermal preference, which may create asymmetrical mating opportunities because of variation in female thermal preference and breadth (F^C^ vs F^W^). The asymmetry of the overlap of males and females at the intermediate developmental temperature could affect sexual selection in populations where Y^M^ and III^M^ both segregate.

At an intermediate developmental temperature (22°C), female thermal preference is bimodal for a reason that we have yet to determine (Figure 3A). This raises the possibility that within populations near the center of the cline (i.e., at a micro-geographic scale), where Y^M^ and III^M^ both segregate (e.g., Hamm & Scott, 2008; Meisel *et al.*, 2016), sexual selection may favor males that can preferentially obtain access to the two different female phenotypes. While differences in thermal preference probably did not evolve in response to sexual selection, these differences do likely have important consequences on the reproductive success of III^M^ and Y^M^ males where they co-occur. III^M^ males may disproportionately benefit from differences in T_pref_ and T_breadth_ between males and females. F^C^ females that prefer colder temperatures have greater T_breadth_ than warm preferring F^W^ females and both male genotypes (Figure 3B), suggesting that F^C^ females occupy a wider range of thermal habitats. Thus, III^M^ males may gain an advantage by having greater access to F^W^ females, as well as occasional access to F^C^ females, in contrast to Y^M^ males who would only be likely to encounter F^C^ females (Figure 4B). This raises the possibility that differences in thermal preference across genotypes and sexes could affect the dynamics of sexual selection.

## Acknowledgements

Tin Nguyen, Chantalle Vincent, and Mandy Cao assisted with experiments. This material is based upon work supported by the National Science Foundation under Grant No. DEB-1845686. Any opinions, findings, and conclusions or recommendations expressed in this material are those of the authors and do not necessarily reflect the views of the National Science Foundation.

## Author Contributions

PD, KA, and RM conceived and designed the study. KA, OH, VS, and JC collected, and KA analyzed, all thermal tolerance data. PD, RP, JT, and AM collected, and PD analyzed, all thermal preference data. PD, KA, and RM wrote the manuscript, and all authors reviewed the manuscript prior to submission.

## Data Accessibility

All data files used for analyses described in this manuscript have been deposited in Dryad (doi:10.5061/dryad.n2z34tmvs). Raw video and image files from thermal preference assays are available from the authors upon request.

## Notes

https://doi.org/10.5061/dryad.n2z34tmvs

### Competing Interest Statement

The authors have declared no competing interest.

### Summary of Updates

This version of the manuscript has been revised to address reviewer comments after submission to a journal.

