## Supplemental Material for "Thermal tolerance and preference are both consistent with the clinal distribution of house fly proto-Y chromosomes"

**Supplementary Materials**

**Supplementary Methods**

*Fly strains and rearing*

We used five nearly isogenic house fly strains, three with $III^M$ males and two with $Y^M$ males, all with a common genetic background. First, CS is an inbred $III^M$ strain that was produced from a mixture of flies collected across the United States (Scott *et al.*, 1996; Hamm *et al.*, 2005). Second, CSrab was created by backcrossing the $III^M$ chromosome from the rspin strain isolated in New York onto the CS background (Shono & Scott, 2003; Son *et al.*, 2019). Third, CSkab was similarly created by backcrossing the $III^M$ chromosome of the KS8S3 strain (Kaufman *et al.*, 2010) onto the CS background. Fourth, IsoCS was created by crossing the $Y^M$ chromosome of a strain collected in Maine onto the CS background (Hamm *et al.*, 2009). Fifth, CSaY was created by backcrossing the $Y^M$ chromosome from the aabys genome reference strain onto the CS background (Scott *et al.*, 2014; Meisel *et al.*, 2015).

We reared each strain at 18°C, 22°C, and 29°C with 12:12-h light:dark photoperiods for two generations. Flies were raised at multiple temperatures because thermal acclimation strongly affects both thermal tolerance (Chown & Terblanche, 2006) and thermal preference (Krstevska & Hoffmann, 1994; Dillon *et al.*, 2009). The strains were maintained with 35 males and 35 females in cages (W17.5 x D17.5 x H17.5 cm) with *ad libitum* supplies of food (1:1 combination of non-fat dry milk and sugar) and water. Females were allowed to lay eggs for 12-24 hours in a standard wheat bran medium (explained in Hamm *et al.*, 2009). We could not collect any eggs from colonies at 18°C, so the 18°C colonies were moved to 22°C in order to collect eggs. Larvae, pupae, and adults from those colonies were subsequently kept at 18°C. We maintained 100-120 larvae per 32 oz container. Within 30 mins of emergence of the third generation, males were separated from females and reared at their developmental temperature (either 18°C, 22°C, or 29°C) until the assays. Females collected within the first half hour of emergence (lighter body color with meconium on their abdomen) are assumed to be unmated.

Flies from each developmental temperature were assayed at different chronological ages to control for differences in the rate of physiological aging across temperatures. For example, one full egg-to-adult generation takes ~1 week at 29°C and ~3 weeks at 18°C. To determine the equivalent physiological age of flies raised at different temperatures, we calculated accumulated degree days ($ADD = T - T_{base}$), where T is the temperature at which the flies were raised (either 18°C, 22°C, or 29°C) and $T_{base}$ is the lower threshold developmental temperature (McMaster & Wilhelm, 1997). $T_{base}$ for house flies is 12.4°C based on previous experiments (Barnard & Geden, 1993; Wang *et al.*, 2018). Total degree days were calculated by multiplying ADD and the total number of days from when eggs were laid until flies were assayed.

For our heat and cold tolerance assays, we used flies 2–3 days post emergence (i.e., eclosion) that developed at 29°C and flies 4-5 days post emergence that developed at 18°C. This is 22–50 total degree days after eclosion. For thermal preference assays, adults raised at 18°C, 22°C or 29°C were assayed at 18–19, 10–11, or 6–7 days after eclosion, respectively. This is approximately 96–115 total degree days after eclosion.

*Thermal tolerance*

To test for heat tolerance, flies were aspirated into an 8 oz paper cup and lightly anaesthetized with $CO_2$. Individual flies were then transferred to a 1.5 ml centrifuge tube, and the opening of the tube was covered with a thin, polyester fabric held in place by a rubber band. The flies were allowed to recover from anaesthesia for at least three hours prior to the assay. Once they recovered, we placed the 1.5 ml tube in a heat block set to 53°C. We chose this temperature because it is the lowest temperature where we observed knockdown in an amount of time that was reasonable for laboratory measurement. Below this temperature, flies were not knocked down in the time period where we could perform the assay. At the time when the tubes were placed in the heat block, all flies clung to the fabric at the top of the tube. The time at which they fell to the bottom of the tube and could not make their way back to the top of the tube was considered the knockdown time.

To test for cold tolerance, lightly anaesthetized flies were transferred to a glass vial (D16 x L150 mm) individually, and the opening of the vial was covered with a thin, polyester fabric held in place by a rubber band. After at least three hours following anesthetization, the vials were placed in a 4°C refrigerator with a transparent door. As in the heat tolerance assay, flies clung onto the fabric at the top of the vial at the beginning of the experiment. Knockdown occured when they fell on their back to the bottom of the vial. However, some flies stuck to the glass surface or the fabric. We therefore gently tapped the assay vial every 2–3 minutes to make sure they were active. The time to knockdown was recorded for each fly. If flies already considered knocked down woke up during tapping of other vials, we waited for them to get knocked down again and noted their final knockdown time.

We performed our experiments in batches, with each batch containing one $III^M$ strain and one $Y^M$ strain. Each batch included 10 males from each combination of genotype ($Y^M$ or $III^M$) and developmental temperature (18°C or 29°C) in a single heat block to measure heat tolerance. Similarly, we assayed 10–15 males per batch from each combination of genotype and developmental temperature to measure cold tolerance. We also included 5 females that developed at each temperature from the same strains as the males in some of the batches as an internal control for both heat and cold tolerance assays. Females from the same strains as the $Y^M$ and $III^M$ males should all have the same (CS) genotype (Meisel *et al.*, 2015). Altogether, we assayed 240 $Y^M$ males, 240 $III^M$ males, and 160 females across 12 batches for heat tolerance, and 450 $Y^M$ males, 450 $III^M$ males, and 220 females over 15 batches for cold tolerance.

We used an analysis of variance (ANOVA) approach to assess the effect of genotype ($Y^M$ vs $III^M$), developmental temperature (18°C or 29°C), and their interaction on time to knockdown in the heat tolerance assays. We did the same for the cold tolerance assays. To do so, we used the lmer() function in the lme4 R package (Bates *et al.*, 2015) to model the effect of genotype (G), developmental temperature (T), and their interaction on knockdown time (KT):

$$KT \sim G + T + G \times T + B + S,$$

with batch (B) and strain (S) treated as random effects. We also constructed another model excluding the interaction term:

$$KT \sim G + T + B + S.$$

We then used a drop in deviance test to compare the fit of the models with and without the interaction term using the anova() function in R.

We also compared heat and cold tolerance between males raised at 22°C and 29°C, using the same approaches as described above. For this comparison, we assayed 140 $Y^M$ males, 140 $III^M$ males across 7 batches for heat tolerance, and 100 $Y^M$ males and 100 $III^M$ males across 5 batches for cold tolerance. We used a drop in deviance test to compare the fit of models with and without an interaction term between genotype and temperature, as described above.

*Thermal preference*

We assessed male thermal preference in the same strains as in the thermal tolerance assays. The sample sizes across groups (strain × developmental temperature combinations) ranged from 19 to 93 adult males. We also conducted thermal preference assays on unmated females from two strains of each male type ($Y^M$: IsoCS and CSaY; $III^M$: CS and CSrab) following the same methods as done with males. The sample sizes across groups (strain × developmental temperature combinations) ranged from 6 to 32 adult females.

We measured thermal preference as the position of individual flies along a thermal gradient, following a slightly modified version of previous protocols (Anderson *et al.*, 2013; Lynch *et al.*, 2018). To create our gradient, we used a 86x20 cm aluminum slab, with one end submerged in a 70°C hot water bath (VWR), and the other end submerged in a styrofoam container filled with crushed ice (Figure S1). Polystyrene insulation was placed underneath the aluminum slab to provide a more consistent gradient. A 10 channel, transparent acrylic cover was created (61x15x1.5 cm) and secured along the gradient with clamps. We drilled six 1 mm holes along the length of each channel to prevent condensation buildup along the gradient. To further prevent condensation buildup, the top of the aluminum slab was lined with filter paper, which was replaced between trials. We built a 3 cm sliding-door acrylic cover that runs the width of the lanes, which was incorporated along the center of the gradient to insert individual flies within each channel. Gradients were allowed to equilibrate for 1 hr before trials began on a given day, generating a temperature range of approximately 17–37°C. All trials were conducted within a 120x55x110 cm PVC frame covered with translucent fabric to homogenize lighting conditions, as house flies have a biased movement towards more intense lighting (Zablocka, 1972). We measured the position of each fly as the linear distance from the left edge of a given lane to the fly within that lane using photographs of the gradient processed in ImageJ (Abràmoff *et al.*, 2004).

To validate the experimental design of our thermal gradient, we first determined the null distribution of movement of adult house flies by recording the behaviors of individuals from two strains (CSkab and IsoCS) at room temperature (22–24°C) with no gradient applied. Individuals (n = 30 per strain) were lightly anesthetized with $CO_2$ for 15 seconds and immediately placed into a gradient channel. Flies were recorded by video camera for 40 minutes, and positions were measured at 5 minute intervals. Fly position followed a unimodal distribution centered at the insertion point, influenced by the fact that anesthetized flies were placed in the center of the gradient and often took several minutes to fully recover from anesthesia and begin movement (Figure S5A).

Next, we followed the methods of Lynch et al. (2018), with slight modifications, to identify a period of low variance in movement from which we obtained behavioral data. Briefly, individuals from CSkab and IsoCS strains (n = 20 flies per strain) were placed onto the gradient for 90 minutes, and individual positions were recorded every minute. We then identified the 10 minute interval where variability in movement began to plateau towards a minimum. This interval corresponded to 25–35 minutes after beginning the assay (Figure S5B). Thus, for all experimental trials, we recorded individual position every minute between 25 and 35 minutes. We excluded data from flies that died before the end of a given trial.

Thermal preference ($T_{pref}$) was estimated based on individual positions along the thermal gradient. Temperatures along the gradient were measured using four thermal sensor probes placed at 12, 24, 36 and 48 cm along the gradient (Figure S1). For each batch, a quadratic function was created based on the known temperatures at the four fixed positions along the gradient. For each individual, we report mean $T_{pref}$ as the average position during the 25–35 minute assay window (measured once per minute). To compare the strength of thermal preference among groups, we measured thermal breadth, $T_{breadth}$ (Carrascal *et al.*, 2016), as the coefficient of variation of individual-level $T_{pref}$ during the 25–35 minute assay window. $T_{breadth}$ provides an estimate of how individuals utilize thermal space within their environment (Slatyer *et al.*, 2013). Choosier individuals show a lower $T_{breadth}$ value and, thus, would be expected to occupy a narrower range of temperatures within a given thermal habitat. As raw $T_{breadth}$ measures did not follow a normal distribution across groups, we used the bestNormalize package in R (v1.2.0) to determine the transformation that most effectively generated a normal distribution of $T_{breadth}$ (Peterson & Cavanaugh, 2019), which was the ordered quantile normalization transformation. We used this normalized value of the coefficient of variation as our $T_{breadth}$ measure.

To determine the effects of developmental temperature (18°C, 22°C, and 29°C), genotype ($Y^M$ and $III^M$), and their interaction on mean $T_{pref}$ across sexes, we created a mixed-effects model using the lme4 package (v1.1) in R (Bates *et al.*, 2015). Developmental temperature, genotype, and their interaction were included as fixed effects, and strain, batch, and lane (L) were included as random effects:

$$T_{pref} \sim G + T + G \times T + B + S + L.$$

We did the same for $T_{breadth}$. Degrees of freedom were estimated using Satterthwaite's formula (Gaylor & Hopper, 1969) using the lmerTest package (v3.1) in R (Kuznetsova *et al.*, 2017). Significance of fixed effects was determined at $p < 0.05$. We then determined whether groups significantly differed in $T_{pref}$ and $T_{breadth}$ using Tukey contrasts with the multcomp package (v1.4) in R (Hothorn *et al.*, 2008).

Within developmental temperature treatments, we used the mclust (v5.4.5) package in R (Scrucca *et al.*, 2016) to determine whether the distribution of individual measures of $T_{pref}$ within a group are best explained by one or multiple normal distributions. We used Bayesian information criterion (BIC) scores to select the best model for a given group. In cases where a given group was best described by a model of multiple normal distributions, we assigned individuals to a given group based on this model. We then tested how $T_{pref}$ and $T_{breadth}$ were each affected by group assignment within a developmental temperature treatment using a mixed-effects model that included strain, batch, and lane as random effects.

**Supplementary Results**

For logistical reasons (skewed male:female ratios, not enough emerging adults, etc.), the sample sizes of different strain x temperature x sex groups were sometimes difficult to control. To account for potential effects caused by this unbalanced design, we have analyzed a down-sampled version of our data and include those results here. Specifically, we used the downsample function of the groupdata2 package in R (Olsen, 2017), and balanced the male and female data sets according to the smallest sample size for a given strain-by-temperature category (n=19 for males, n=6 for females). We conducted mixed-model analysis of variance using the same models as explained in the manuscript, then extracted *p*-values. We repeated this process of downsampling and extracting *p*-values 1,000 times and report the mean *p*-values of each fixed effect here.

In general, downsampling to the smallest sample sizes in both male and female data sets does not greatly influence our findings. For male $T_{pref}$, we still observe significant main effects of genotype ($p = 0.019$) and developmental temperature ($p < 1.0 \times 10^{-6}$). The interaction effect of genotype x temperature becomes marginally non-significant ($p = 0.052$). However, we still find that $III^M$ and $Y^M$ males exhibit significantly different thermal preferences when developed at the intermediate temperature of 22℃ (Tukey's post-hoc, $p < 1.0 \times 10^{-6}$), but not when developed at 18℃ ($p = 0.72$) or 29℃ ($p = 0.37$). For female $T_{pref}$, we still observe only a significant main effect of temperature on female $T_{pref}$ ($p = 4.3 \times 10^{-4}$), and no significant effects of genotype or the genotype x temperature interaction (both $p > 0.3$).

Lastly, to estimate effects of male and female group assignments ($III^M$ and $Y^M$ males, $F^C$ and $F^W$ females) on $T_{breadth}$, we balanced the combined male and female data set of preference behaviors recorded at 22℃ according to the smallest sample size for a given strain x sex category (n=19). We conducted mixed-model analysis of variance as previously described, followed by

Tukey's post-hoc tests, and we extracted *p*-values for pairwise comparisons of $T_{breadth}$ between groups. We repeated this process 1,000 times and report the mean *p*-values here. Similar to our manuscript results, we found that cold-preferring females ($F^{C}$) show significantly higher $T_{breadth}$ values than $F^{W}$ females ($p = 7.6 \times 10^{-4}$), $III^{M}$ males ($p = 1.4 \times 10^{-4}$), and $Y^{M}$ males ($p = 6.3 \times 10^{-4}$). All other pairwise comparisons were not significantly different (all $p > 0.8$).

**Supplementary Figures**

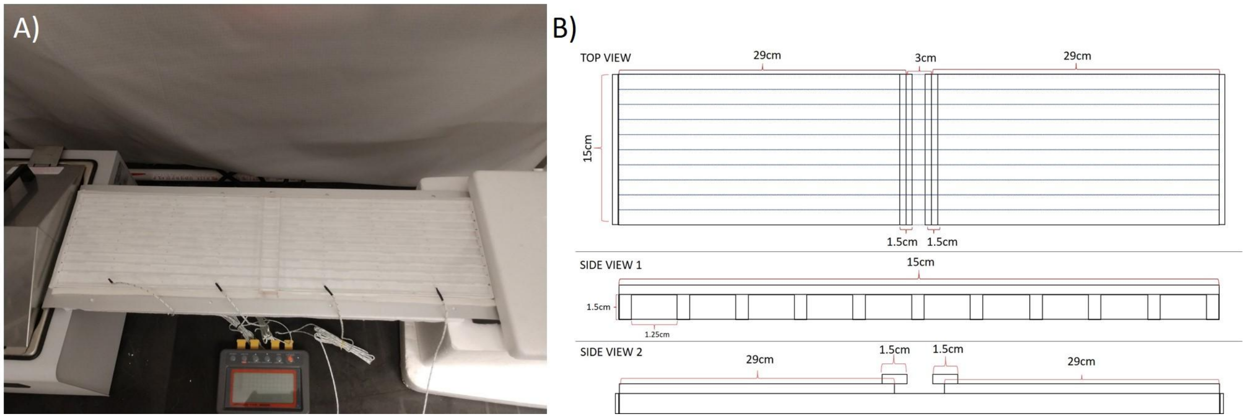


**Figure S1**- Thermal gradient design. (**A**) Photograph of thermal gradient on aluminum slab, with ends inserted into hot water bath (left) and crushed ice container (right). Four temperature probes are inserted into 1 mm holes evenly distributed along the gradient. (**B**) Schematic diagram for construction of clear acrylic lid. Along each lane, six 1 mm holes are drilled to prevent condensation buildup within lanes.

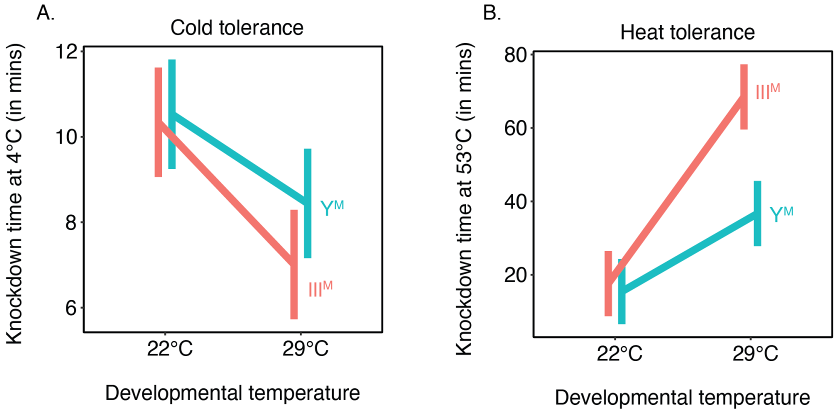


**Figure S2**- Cold tolerance (**A**) and heat tolerance (**B**) in males raised at 22°C and 29°C. Each data point is an estimated mean. Error bars represent standard error.

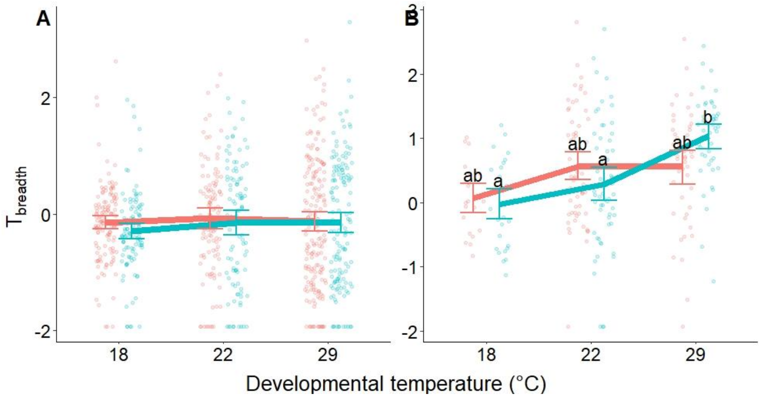


**Figure S3** - Thermal breadth of (**A**) male and (**B**) female house flies according to genotype ($III^M$ = salmon points and line, $Y^M$ = turquoise points and line) and developmental temperature. Each point depicts the thermal breadth for an individual fly, with lines and error bars denoting means within group and standard errors of the mean, respectively. In males, no groups were significantly different in $T_{breadth}$. In females, significant differences between groups are denoted by letters, with differing letters highlighting significantly different $T_{breadth}$ (Tukey's *post hoc* test, $p < 0.05$).

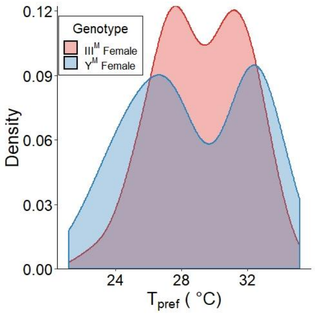


**Figure S4**- Distributions of individual-level mean thermal preferences of females according to male genotype in the strain (females from strains with $III^M$ males are shown in red, females from strains with $Y^M$ males are shown in blue). The Y-axis represents relative density of data points and is analogous to frequency of data points for a given $T_{pref}$ value.

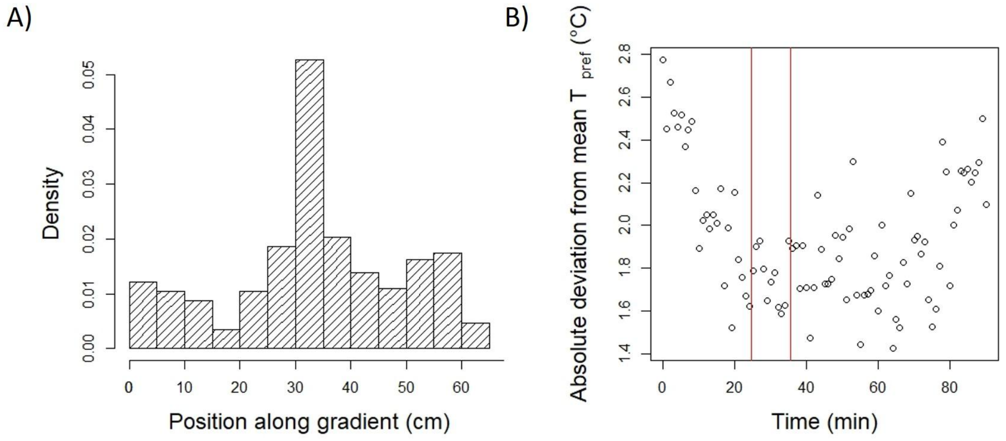


**Figure S5**- Summary of validation experiments for thermal preference assays. (**A**) Density histogram of individual positions along thermal gradient (in cm, from left to right). (**B**) The absolute deviation from individuals' mean preference, at every minute across the 90-minute pilot trials. Red lines indicate the 25–35 minute interval used for subsequent thermal preference assays.

### Supplementary Tables

**Table S1** - Summary statistics showing properties of mixture models fit to $T_{pref}$ values for $III^M$ male, $Y^M$ male, and female house flies. The BIC values and summary statistics of the best fit models are shown in bold. df - degrees of freedom.

| Group | Number of Distributions | BIC (equal $\sigma^2$) | BIC (unequal $\sigma^2$) | df | Distribution | % Flies in Distribution | Mean (℃) | $\sigma^2$ (℃) |
|---|---|---|---|---|---|---|---|---|
| $III^M$ ♂ | 1 | -578.6 | -578.6 | 2 | 1 of 1 | 100 | 30.1 | 10.3 |
| | 2 | -570.3 | **-556.8** | **5** | **1 of 2** | **60** | **28.7** | **10.4** |
| | | | | | **2 of 2** | **40** | **32.6** | **0.42** |
| | 3 | -579.7 | -569.6 | 8 | 1 of 3 | 32.7 | 27.2 | 8.76 |
| | | | | | 2 of 3 | 30 | 31 | 5.56 |
| | | | | | 3 of 3 | 37.3 | 32.5 | 0.26 |
| $Y^M$ ♂ | 1 | **-527.4** | **-527.4** | **2** | **1 of 1** | **100** | **26.9** | **12.24** |
| | 2 | -536.2 | -540.8 | 4 | 1 of 2 | 49.5 | 24.8 | 7.76 |
| | | | | | 2 of 2 | 50.5 | 29.1 | 7.76 |
| | 3 | -545 | -554.2 | 6 | 1 of 3 | 21.6 | 23.2 | 5.02 |
| | | | | | 2 of 3 | 59.8 | 27.1 | 5.02 |
| | | | | | 3 of 3 | 18.6 | 31.1 | 5.02 |
| ♀ | 1 | -619.6 | -619.6 | 2 | 1 of 1 | 100 | 28.9 | 10.3 |
| | 2 | -614.9 | **-610.2** | **5** | **1 of 2** | **59.3** | **26.9** | **4.93** |
| | | | | | **2 of 2** | **40.7** | **32.2** | **1.28** |
| | 3 | -611.5 | -618.1 | 6 | 1 of 3 | 12.7 | 24.0 | 1.55 |
| | | | | | 2 of 3 | 42.4 | 27.3 | 1.55 |
| | | | | | 3 of 3 | 44.9 | 32.0 | 1.55 |
